# Synapsin-caveolin-1 mitigates cognitive deficits and neurodegeneration in Alzheimer’s disease mice

**DOI:** 10.1101/2020.07.24.220129

**Authors:** Shanshan Wang, Joseph S. Leem, Sonia Podvin, Vivian Hook, Natalia Kleschevnikov, Paul Savchenko, Mehul Dhanani, Kimberly Zhou, Isabella C. Kelly, Tong Zhang, Atsushi Miyanohara, Alexander Kleschevnikov, Steve L. Wagner, John Q. Trojanowski, David M. Roth, Hemal H. Patel, Piyush M. Patel, Brian P. Head

## Abstract

AD presents with severe neurodegeneration which leads to cognitive deficits and dementia. Identifying the molecular signals that attenuate neurodegeneration in AD may be exploited as therapeutic targets. This study revealed that transgenic AD mice (PSAPP) exhibit decreased caveolin-1 (Cav-1), a membrane/lipid raft (MLR) scaffolding protein that organizes synaptic signaling components. Subcellularly, Cav-1 and full length (fl)-TrkB were significantly decreased in MLRs. We thus developed an in vivo gene therapy that re-expresses neuronal-targeted Cav-1 using the synapsin promoter (*SynCav1*). While AD mice showed significant learning and memory deficits at 9 and 11 months, AD mice that received hippocampal *SynCav1* (AD-*SynCav1*) maintained normal learning and memory at 9 and 11 months respectively. Furthermore, AD-*SynCav1* mice showed preserved hippocampal MLR-localized fl-TrkB, synaptic ultrastructure, dendritic arborization and axonal myelin content, all of which occurred independent of reducing amyloid deposit and astrogliosis. Thus, *SynCav1* demonstrates translational potential to treat AD by delaying neurodegeneration.

**Summary:** Transgenic PSAPP mice exhibit decreased hippocampal expression of the membrane lipid raft (MLR) scaffolding protein caveolin-1. Synapsin-promoted re-expression of Cav-1 (termed *SynCav1*) mitigated neuropathology and cognitive deficits. *SynCav1* gene therapy has the potential to treat AD and other forms of neurodegeneration.

## Introduction

Alzheimer’s disease (AD) is a devastating neurodegenerative condition and the most common cause of dementia. Although AD is characterized by abundant amyloid-b (Ab) plaques and neurofibrillary tau tangles, disrupted synaptic signaling and loss of synapses and neurons are more closely correlated to cognitive deficits. Unfortunately, efforts to combat the disease have been hindered by the growing list of failed clinical trials with interventions focused on reducing amyloid-related pathology, thus causing many to reconsider the amyloid hypothesis as the primary cause of AD(Doody et al., 2014; Hardy & Higgins, 1992; Honig et al., 2018; Salloway et al., 2014). Failed clinical efforts which focused on reducing Ab suggest that removal of toxic amyloid species alone is not sufficient to evoke functional neuronal and synaptic plasticity in the neurodegenerative brain. Gene therapies which target neuroprotection and resilience have immense potential to treat individuals afflicted with AD or other forms of neurodegeneration of different or unknown etiology(Bouaita et al., 2012; Eggers et al., 2019).

Cellular plasma membranes contain discrete microdomains enriched in synaptic signaling components and cholesterol termed membrane/lipid rafts (MLRs). Evidence shows that toxic Ab species aggregate with membrane cholesterol and other lipids resulting in damage or loss of MLR-related signaling proteins, ultimately leading to synaptic loss and cognitive dysfunction in AD(Fabelo et al., 2014; Morgado & Garvey, 2015). The disruption of MLRs and decreased MLR-associated scaffolding proteins may in part explain for the lack of efficacy in clinical trials that attempted to restore function through delivery of exogenous neurotrophins to AD patients using *AAV2-NGF* gene delivery(Rafii et al., 2018). One potential alternative to treat AD and other neurodegenerative conditions is the use of gene therapies that target neuroprotective mechanisms via restoration of MLR-localized functional NTRs(Combs, Kneynsberg, & Kanaan, 2016; Deverman, Ravina, Bankiewicz, Paul, & Sah, 2018). Enhancing MLR-localization of NTRs may augment pro-survival and pro-growth signaling, enhance structural and functional neuroplasticity, or even increase the efficacy of exogenous or already present endogenous growth factors, which in turn may restore cognitive function in AD. One such gene therapy candidate is caveolin-1 (Cav-1), a MLR scaffolding protein that organizes NTRs (Trk) and neurotransmitter receptor signaling complexes in MLRs(Egawa et al., 2017; Head et al., 2011; Mandyam et al., 2017). Both pre-clinical and clinical findings revealed that Cav-1 and Cav-1 associated signaling complexes (NTRs and neurotransmitter receptors) were decreased in degenerating neurons in AD, chronic traumatic encephalopathy (CTE), and amyotrophic lateral sclerosis (ALS)(Flis et al., 2018; Mufson et al., 2018; Tiernan et al., 2018; Yuan et al., 2017). In contrast, we previously showed that neuron-targeted Cav-1 over-expression (i.e., synapsin-promoted Cav-1 or *SynCav1*) augmented agonist-mediated synaptic signaling (e.g., NTRs, neurotransmitter receptors) and dendritic arborization *in vitro*(Head et al., 2011), preserved MLR-localized TrkB and extended life span in ALS mice(Sawada et al., 2019), suggesting that alterations in Cav-1 expression affects pro-survival signaling and neuroprotection in neurodegenerative conditions.

The present study tested whether *SynCav1* gene delivery to AD mice could restore MLR-localized TrkB, preserve neuronal and synaptic plasticity, and improve higher brain function. These findings are the first to demonstrate that a one-time hippocampal delivery of *AAV-SynCav1* to 3-month old AD mice preserved hippocampal learning and memory at 9 and 11 months. Moreover, *SynCav1* preserved MLR-localization of TrkB, hippocampal dendritic arborization, synaptic ultrastructure, and axon myelin content. Thus, *SynCav1* gene therapy is an attractive approach to preserve brain plasticity and improve higher brain function in AD and potentially in other forms of neurodegeneration resulting from unknown etiology.

## Results

### *SynCav1* preserves fear learning and memory in 9 and 11 m AD mice

We first tested whether direct hippocampal delivery of AAV9 vectors would affect cognitive performance. No significant difference was observed in open field or fear conditioning behavioral tests between naïve WT mice versus WT-*SynRFP* (Supplementary Fig. 3). Because there was no effect from AAV vectors on behavior, we did not include a naïve WT control group (i.e., non-AAV) in this study. Previous work from our group demonstrated augmented functional neuroplasticity and improved hippocampal-dependent memory in WT mice that received AAV9-*SynCav1*(Egawa et al., 2018; Mandyam et al., 2017), therefore in order to reduce experimental redundancy the WT-*SynCav1* group was not included in the present study. To test the hypothesis that *SynCav1* can improve brain function in a neurodegenerative AD mouse model, the current study utilized the following 3 groups to test this hypothesis: WT-*SynRFP*, AD-*SynRFP* and AD-*SynCav1*.

Open field at 9 and 11 months (m) revealed no general motor deficits nor anxiety-like behavior among the 3 groups (Supplementary Fig. 4). At 9 m, AD-*SynRFP* mice exhibited a significant reduction in fear learning acquisition on day 1 compared to WT-*SynRFP* (Fig. 1E). In contrast, 9 m AD-*SynCav1* mice exhibited preserved fear learning on day 1 (no difference vs WT-*SynRFP*). No significant difference in contextual (Day 2) or cued memory recall (Day 3) were observed among the groups. These results demonstrate that *SynCav1* gene delivery preserves fear learning in mild symptomatic AD mice.

**Fig. 1.**
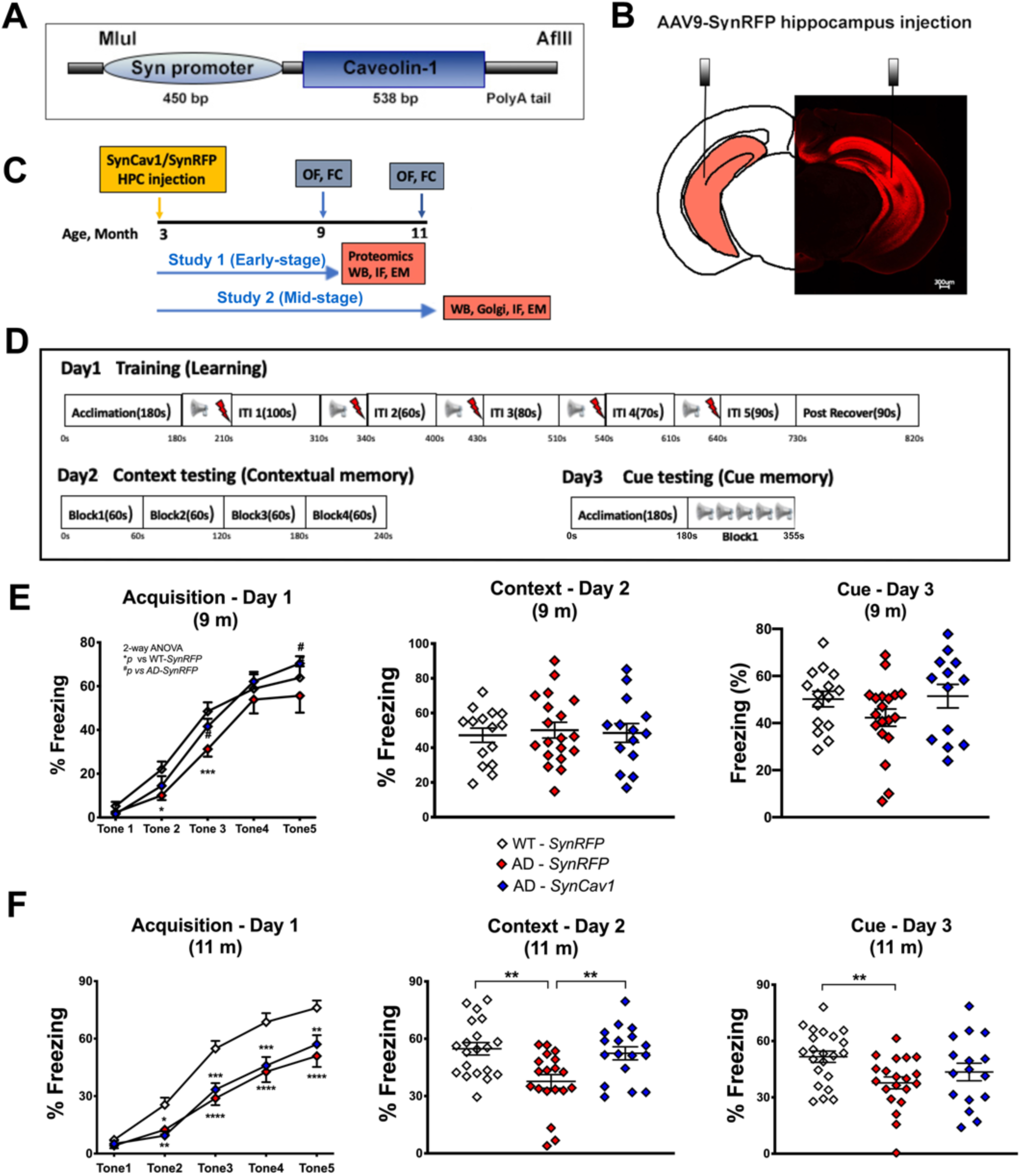
*SynCav1* gene delivery preserves learning and memory in 9 and 11 month-old APPSwePS1d9 (AD) mice. (**A**) Viral structure of AAV9-*SynCav1*. (**B**) Illustration (left) and microscopy of hippocampal RFP. (**C**) Schematic of experimental design in WT and APPSwePS1d9 (AD) mice. Blue arrows and rectangular boxes indicate open field and fear conditioning tests for separate 9 and 11 m cohorts; orange arrows and rectangular boxes indicate postmortem biochemical assays. (**D**) Schematic of fear conditioning protocol. Fear conditioning test showed that at 9 m (**E**), AD-*SynRFP* mice only exhibited reduction in fear learning acquisition on day 1 compared to WT-*SynRFP* with normal contextual memory recall on day 2. At 11 m (**F**), both AD groups exhibited a significant reduction in learning on day 1 compared to WT-*SynRFP*. On day 2, AD-*SynRFP* mice exhibited a significant reduction in contextual memory recall versus WT-*SynRFP* while AD-*SynCav1* exhibited normal contextual memory recall. Data are presented as percent (%) freezing mean ± SEM. Data were analyzed using two-way analysis of variance (ANOVA) followed by Fisher’s LSD multiple comparisons tests (Day 1) or Students *t* test (Day 2 and 3). *n* = 14-19 animals per group for 9 m; *n* = 17-22 per group for 11 m. Significance was assumed when *p* < 0.05. \**p* < 0.05, \*\**p* < 0.01, \*\*\**p* < 0.001, \*\*\*\**p* < 0.0001.

At 11 m, both AD groups exhibited a significant reduction in learning on day 1 compared to WT-*SynRFP* (Fig. 1F). On day 2, AD-*SynRFP* mice exhibited a significant reduction in contextual memory recall versus WT-*SynRFP* and versus AD-*SynCav1*, indicating hippocampal memory deficits. There was no significant difference between AD-*SynCav1* versus WT-*SynRFP* on day 2 demonstrating preserved hippocampal-dependent memory recall in AD mice that received *SynCav1*. On day 3, there was a significant decrease in AD-*SynRFP* percent freezing versus WT-*SynRFP*; no difference was observed between AD-*SynCav1* versus WT-*SynRFP*. These data demonstrate direct evidence that *SynCav1* gene delivery preserves hippocampal memory in symptomatic AD mice.

### *SynCav1* preserves MLR-localization of Cav-1 and TrkB in hippocampal tissue from symptomatic AD mice

Immunofluorescence (IF) of 9 m AD-*SynRFP* mice showed decreased hippocampal Cav-1 expression, which was preserved in AD-*SynCav1* (Fig. 2A). Co-staining of the dendritic marker MAP2 with Cav-1 showed decreased MAP2 expression in the hippocampal CA1 subfield and cortex in AD-*SynRFP*, similar to that observed in postmortem human brains of patients diagnosed with CTE, tauopathy, and AD(Mufson et al., 2018; Tiernan et al., 2018). In contrast, AD-*SynCav1* mice displayed increased hippocampal (Fig. 2B) and cortical (Fig. 2C) MAP2 expression. Immunoblot (IB) of hippocampal homogenates confirmed decreased Cav-1 expression in AD-*SynRFP* at 9 m (Fig. 2D, E). Although no significant difference was found between WT-*SynRFP* versus AD-*SynRFP*, AD-*SynCav1* mice exhibited a significant increase in full-length TrkB (fl-TrkB) versus AD-*SynRFP* mice at 9 m. *SynCav1* had no effect on APP expression. IB of buoyant MLR fractions revealed a significant decrease in Cav-1 and fl-TrkB in AD-*SynRFP* versus WT-*SynRFP*. In contrast, MLRs from AD-*SynCav1* mice exhibited a significant increase in Cav-1 and fl-TrkB (versus AD-*SynRFP* mice (Fig. 2D, E). No change in MLR-associated synaptobrevin expression (data not shown) was observed among the 3 groups, thus confirming the MLR changes were specific to Cav-1 and fl-TrkB. The decreased MLR-localized fl-TrkB expression in AD mice indicates a subcellular alteration which may in part contribute to neuronal dysfunction in AD. Furthermore, the subcellular alteration in TrkB in the hippocampus from AD mice was prevented with *SynCav1*.

**Fig. 2.**
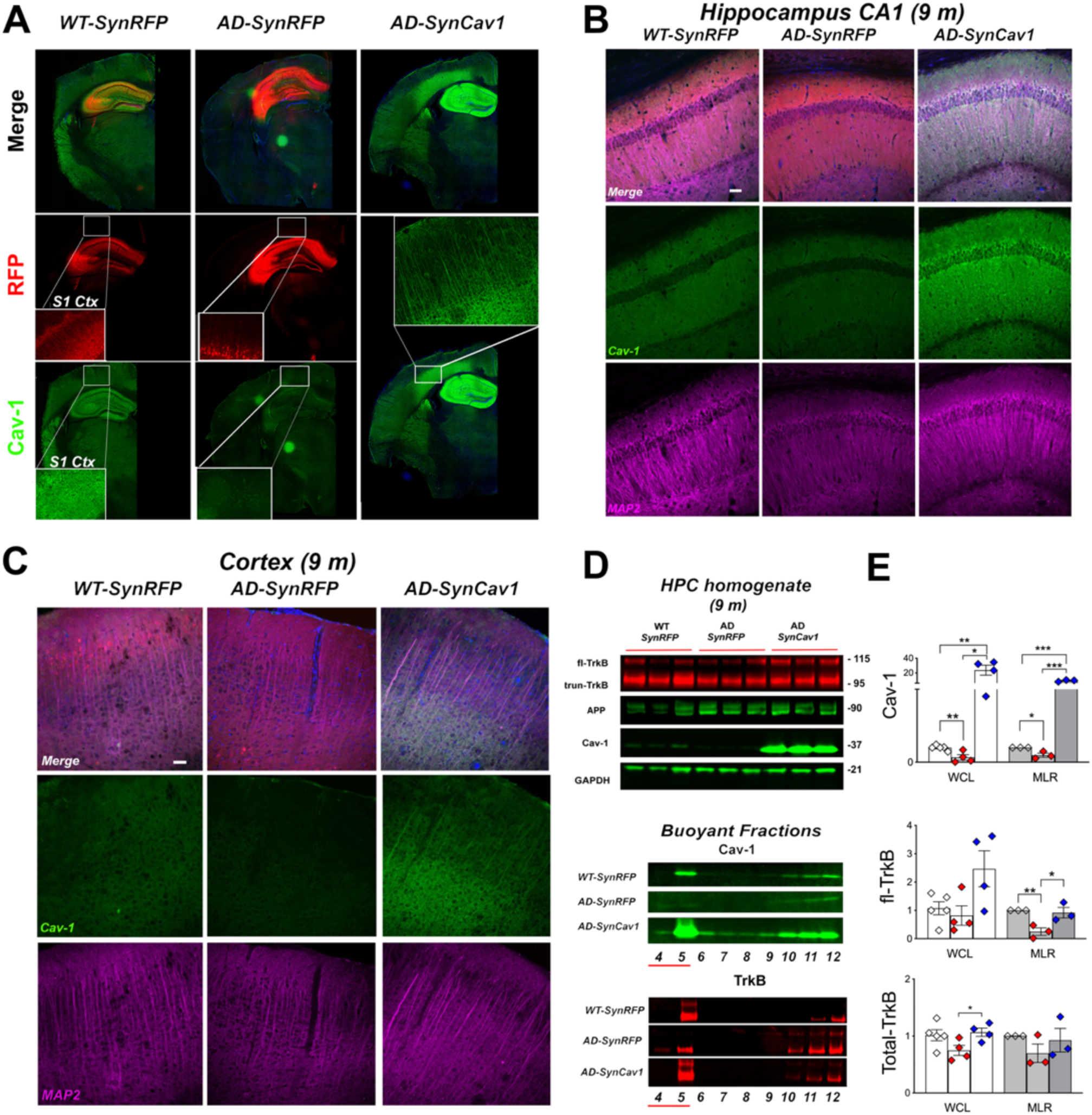
*SynCav1* gene delivery preserves hippocampal Cav-1 expression and MLR-localized fl-TrkB in 9 m APPSwePS1d9 (AD) mice. (**A**) Immunofluorescence microscopy revealed significantly decreased Cav-1 in 9 m AD mice hippocampus. (**B-C**) Co-staining for Cav-1 and dendritic marker MAP2 showed that *SynCav1* gene delivery preserved MAP2 expression in dendritic processes in the hippocampal CA1 subfields CA and cortex in 9-month old AD mice. Scale bar = 50 um. IB (**D**-**E**) of 9 m hippocampal homogenates and buoyant fractions (i.e., MLRs) revealed decreased Cav-1 and full-length TrkB (fl-TrkB) expression in AD-*SynRFP* mice vs WT-*SynRFP* mice. *SynCav1* preserved hippocampal MLR localized fl-TrkB expression in AD mice. Fractions were generated from equal protein (0.5 ug/ul). Data present as mean ± SEM, *n* = 3-4 animals per group. Data were analyzed using Student *t* test. Significance was assumed when *p* < 0.05. \**p* < 0.05, \*\**p* < 0.01, \*\*\**p* < 0.001.

### Proteomics analysis of MLRs reveals decreased expression of neurodegenerative genes in AD-*SynCav1* in mice

Tandem mass spectrometry (MS/MS) assessed proteins in MLRs from WT-*SynRFP*, AD-*SynRFP* and AD-*SynCav1*. Quantifiable protein groups were first evaluated for significant expression differences between WT-*SynRFP* and AD-*SynRFP* (Fig. 3). Of 2417 quantifiable proteins, 65 were significantly upregulated, with mean log_2_ (AD-*SynRFP* /WT-*SynRFP*) > 0, and 52 significantly downregulated, with mean log_2_ (AD-*SynRFP* /WT-*SynRFP*) < 0 (Fig. 3A). Mean expression differences of significantly regulated proteins were evaluated by complete Euclidean hierarchical clustering to identify protein groups of mild, moderate, or strong up or down-regulation (Fig. 3B). Cav-1 was also significantly downregulated in MLRs from AD-*SynRFP* hippocampi (denoted by red box in Bacterial Invasion of epithelial Pathways graph), which is consistent with the IB results shown in Fig. 2C. Clustering revealed Shisa9 (an AMPAR regulatory protein) was the most strongly downregulated protein in AD-*SynRFP* compared to WT-*SynRFP* (log_2_ = -5.9). All significantly regulated quantifiable protein groups together evaluated for significant gene ontology (GO) functional enrichment using STRING-db (www.string-db.org) (Fig. 3C). The most significant GO terms identified were Bacterial Invasion of Epithelial Cells, Ribosome, Ferroptosis, Mineral Absorption, and Adherens Junction. GO genes for Bacterial Invasion of Epithelia contained Cav-1, which was also significantly downregulated in AD-*SynRFP* yet preserved in AD-*SynCav1*.

**Fig. 3.**
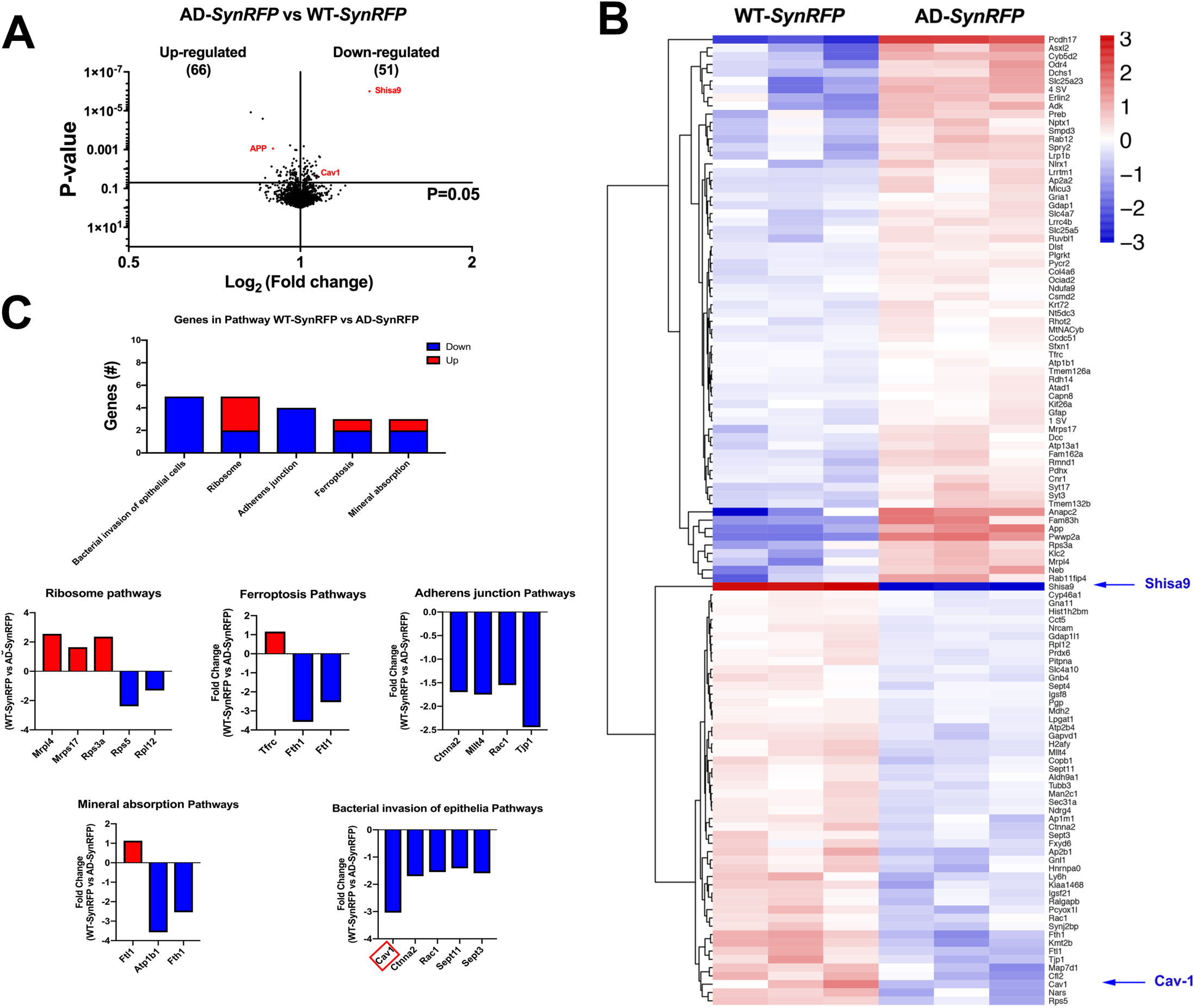
Proteomics reveals altered MLR-associated proteins in AD-*SynRFP* mice at 9 m. Volcano plot of 9 m MLRs between WT-*SynRFP* and identified 117 genes that were differentially expressed (65 upregulated and 52 down regulated) (**A**). Mean expression differences of significantly regulated proteins were evaluated by Euclidean hierarchical clustering (**B**). Gene ontology (GO) functional enrichment using STRING-db (**C**). Clustering revealed Shisa9 (an AMPAR regulatory protein) was the most strongly downregulated protein in AD-*SynRFP* compared to WT-*SynRFP* (log_2_ = -5.9). Cav-1 was also significantly downregulated in MLRs from AD-*SynRFP* hippocampi. All significantly regulated quantifiable protein groups together evaluated for significant gene ontology (GO) functional enrichment using STRING-db (www.string-db.org). The most significant GO terms identified were Bacterial Invasion of Epithelial Cells, Ribosome, Ferroptosis, Mineral Absorption, and Adherens Junction. All proteins in this GO term and Adherens Junction were downregulated in AD-*SynRFP* versus WT-*SynRFP*. The significantly different expression levels between comparison groups were determined by two-tailed Student *t* test, *n* = 3 animals per group. Significance was assumed when \**p* < 0.05.

We next compared AD-*SynRFP* versus AD-*SynCav1* (Fig. 4). Of 2417 proteins, 168 were significantly altered, with 80 significantly upregulated and 88 significantly downregulated (Fig. 4A,B). Cav-1 was the most upregulated protein (log2 = 7.8) due to its overexpression with *SynCav1*. Shisa9 was the second most upregulated protein (log2 = 5.9), as confirmed by IB (Supplementary Fig.5). Notably, several downregulated proteins in MLRs from AD-*SynRFP* (ap2b1, cofilin-2, cct-5, FXYD, Kiaa1468, MAN2C1, NDRG4, synaptojanin2, Prdx6) were significantly upregulated in MLRs from AD-*SynCav1*. Furthermore, several genes implicated in neurodegenerative disease pathways (e.g., AD, Parkinson’s, Huntington’s) were significantly down regulated in MLRs from AD-*SynCav1* when compared to AD-*SynRFP* (Fig. 4C, graphs in blue rectangle inset). Bacterial invasion of epithelia graph shows significant increase in Cav-1 (Fig. 4B, red rectangle). These results show that *SynCav1* decreases MLR expression of genes implicated in neurodegenerative pathways.

**Fig. 4.**
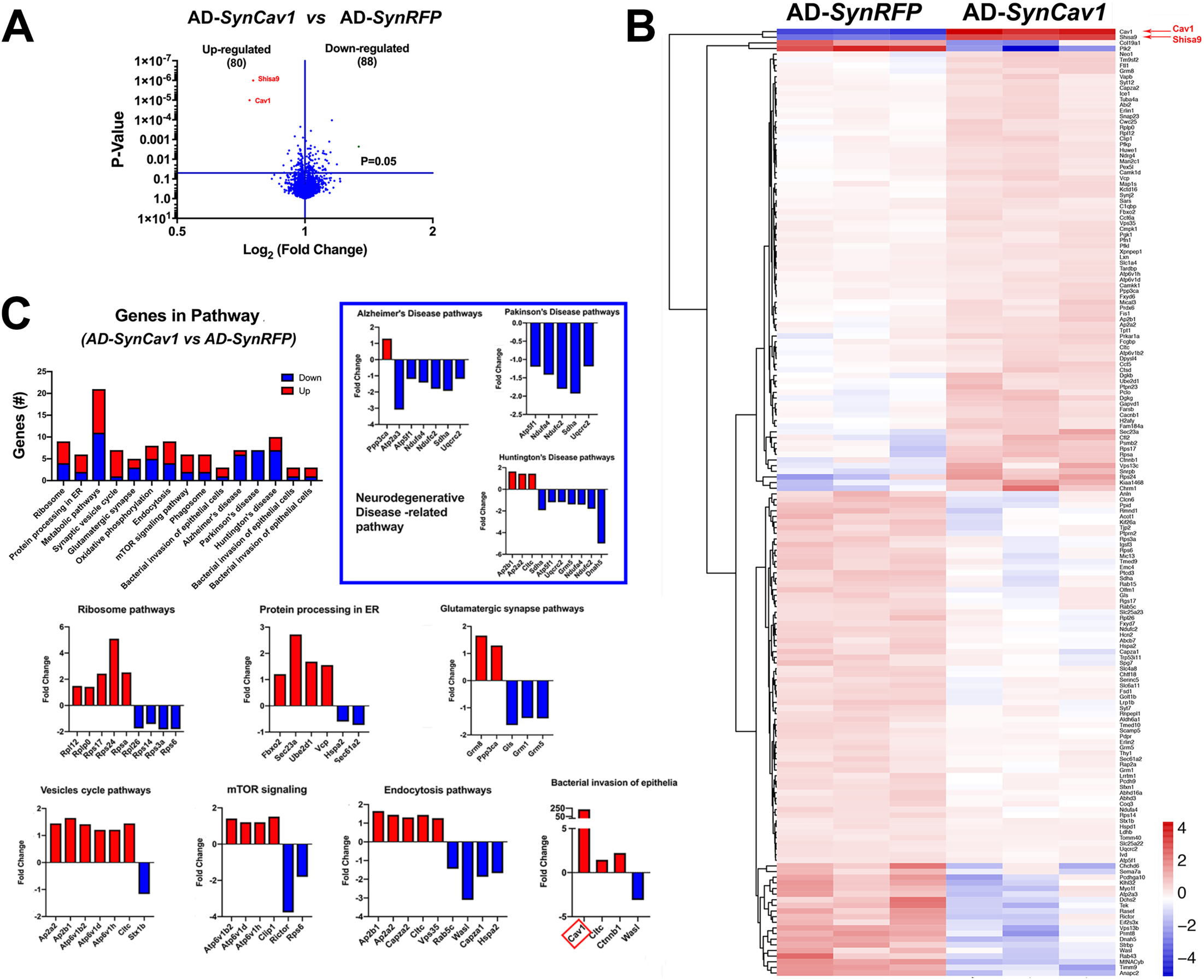
Proteomics reveals that *SynCav1* decreases expression of neurodegenerative genes in MLR fractions from 9 m AD mice. Volcano plot of 9 m MLRs between AD-*SynRFP* and AD-*SynCav1* identified 168 genes that were differentially expressed (80 upregulated and 88 down regulated) (**A**). Mean expression differences of significantly regulated proteins were evaluated by Euclidean hierarchical clustering (**B**). Cav-1 was the most upregulated protein (log2 = 7.8) due to its overexpression with *SynCav1*. Shisa9 was the second most upregulated protein (log2 = 5.9). Notably, several downregulated proteins in MLRs from AD-*SynRFP* (ap2b1, cofilin-2, cct-5, FXYD, Kiaa1468, MAN2C1, NDRG4, synaptojanin2, Prdx6) were significantly upregulated in MLRs from AD-*SynCav1*. Gene ontology (GO) functional enrichment using STRING-db demonstrated that several genes implicated in neurodegenerative disease pathways (e.g., AD, Parkinson’s, Huntington’s) were significantly down regulated in MLRs from AD-*SynCav1* when compared to AD-*SynRFP* (**C**). Red rectangular box in Bacterial Invasion of Epithelia Pathways in **c** denotes increased Cav-1. The significantly different expression levels between comparison groups were determined by two-tailed Student *t* test, *n* = 3 animals per group. Significance was assumed when \**p* < 0.05.

### AD-*SynCav1* mice exhibit preserved dendritic arborization and synaptic ultrastructure in CA1 hippocampal neurons in 11 m AD mice

Changes in hippocampal dendritic arborization are necessary for memory. Golgi-Cox staining (Fig. 5A) revealed a significant reduction in CA1 apical and basal dendritic arborization, apical dendritic soma to tip distance and dendritic spines in AD-*SynRFP* mice compared to WT-*SynRFP*. AD-*SynCav1* showed preserved apical dendritic arborization from 70 to 280 um distance from the soma (Fig. 5B), preserved CA1 basal dendritic arborization (Fig. 5B, *right graph*), greater apical dendrite soma to tip distance (Fig. 5C), with no difference in basal dendrite soma to tip distance, and increased number of CA1 dendritic spines (Fig. 5D). These findings indicate that *SynCav1* preserves structural neuroplasticity in the hippocampus of AD mice, a structural preservation that may in part explain for the preserved hippocampal-dependent memory measured in Fig. 1E.

**Fig. 5.**
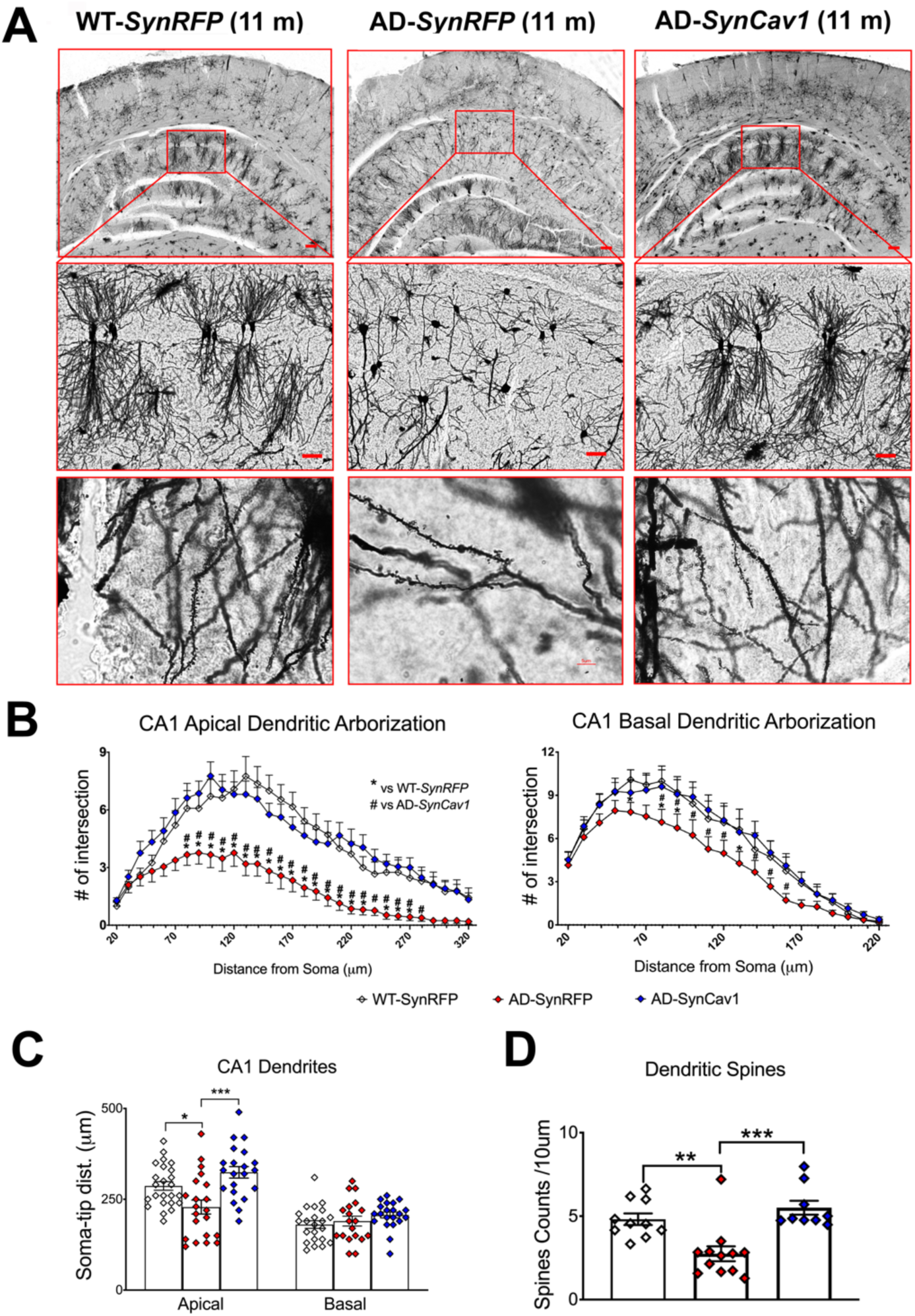
*SynCav1* gene delivery preserves dendritic arborization in CA1 hippocampal neurons from 11 m AD mice. Golgi-Cox images of *cornu ammonis* (CA1) pyramidal neurons (Top row scale bar = 200 um; middle row images scale bar = 50 um) (**A**). Quantification revealed a significant reduction in CA1 apical and basal dendritic arborization in AD-*SynRFP* mice compared to WT-*SynRFP* and AD-*SynCav1* mice. AD-*SynCav1* showed preserved apical dendritic arborization from 70 to 280 um distance from the soma and basal dendritic arborization from 80 to 150 um distance (**B**). AD-*SynRFP* mice also showed decreased apical dendrite soma to tip distance, which was preserved by *SynCav1* (**C**). No difference in basal dendrite soma to tip distance (**C**). Further analysis revealed decreased CA1 spine number in AD-*SynRFP* mice and preserved dendritic spines in AD-*SynCav1* (**D**). Data are presented as mean ± SEM, *n* = 3 animals in each group consisting of 8-12 neurons from each animal. Data were analyzed using Student *t* test. Significance was assumed when *p* < 0.05. \**p* < 0.05, \*\**p* < 0.01, \*\*\**p* < 0.001.

### *SynCav1* preserves infrapyramidal mossy fiber area and CA3 Schaffer axon myelination in the hippocampus of AD mice

Proper hippocampal circuitry (Fig. 6A) is essential for cognitive function. The mossy fiber pathway, which consists of unmyelinated axons projecting from the dentate gyrus to CA3 pyramidal cells, strongly correlates with spatial learning(Pearn et al., 2018). Because AD-*SynRFP* mice showed a significant reduction in hippocampal-dependent memory recall at 11 m, we measured infrapyramidal mossy fiber (IPM) and suprapyramidal mossy fiber (SPM) area by synaptoporin staining (Fig. 6B)(Pearn et al., 2018). As shown in Fig. 6B, AD-*SynRFP* mice exhibited significantly reduced IPM area compared to WT-*SynRFP*, while the AD-*SynCav1* group showed significantly greater IPM area compared to AD-*SynRFP*, with no significant difference in SPM area detected (Fig. 6C-6E).

**Fig. 6.**
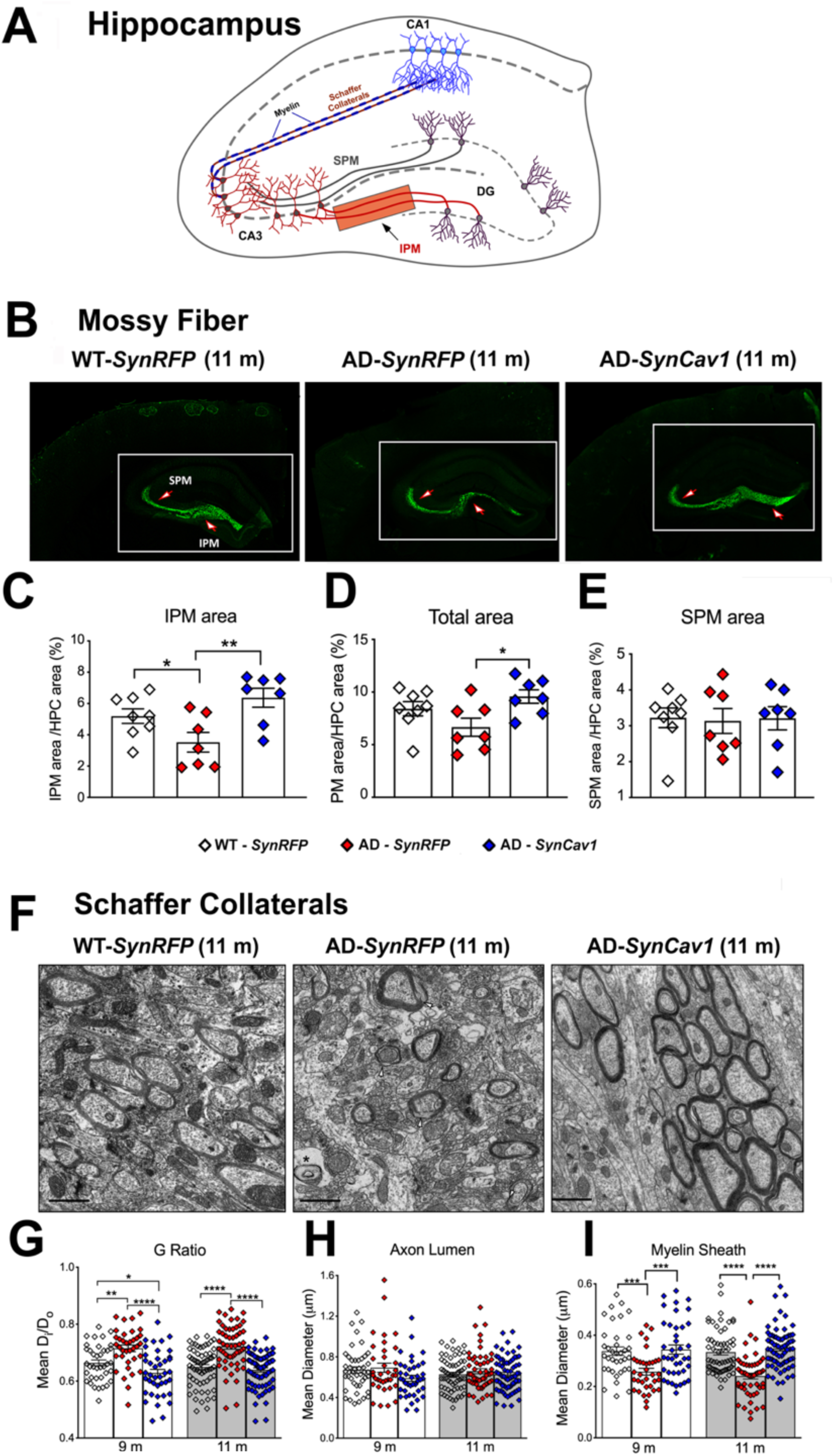
*SynCav1* gene delivery preserves axonal structure and myelin content in the hippocampus of 9 m and 11 m AD mice. Illustration of hippocampal pathways (**A**). Infrapyramidal and suprapyramidal mossy fiber structures at 11 m (*n* = 3-4 animals per group consisting of 2-3 images per animal) (**B**). AD-*SynRFP* mice exhibited significantly reduced IPM area (**C, D**) compared to WT-*SynRFP*, while the AD-*SynCav1* group showed significantly greater IPM area compared to AD-*SynRFP*, with no significant difference in SPM area detected (**E**). EM images of myelinated CA3 Schaffer collateral axons (**F**). G-ratio analysis of CA3 Schaffer collateral axons (axon lumen diameter (D_i_)/fiber (axon lumen + myelin) diameter (D_o_)) revealed that the G-ratio was increased in AD-*SynRFP* mice at 9 m versus WT-*SynRFP* (**G**), an alteration due to reduced myelin sheath diameter (**I**), rather than increased axon lumen diameter (**H**). AD-*SynCav1* mice exhibited significantly decreased G-ratio compare to AD-*SynRFP* due to increased myelin sheath diameter. Further analysis of 11 m revealed similar increase in the G-ratio of AD-*SynRFP* mice (**G**) compared to WT-*SynRFP*, while AD-*SynCav1* mice exhibited a significant decrease in G-ratio verse AD-*SynRFP*, the G-ratio change was due to altered myelin sheath thickness (**I**) rather than axon lumen diameter (**H**). Micrographs were captured at 11000x Magnification, scale bar = 500 nm. Data are presented as mean ± SEM, *n* = 3-5 animals per group consisting of 10 electron micrographs images per animal. Data were analyzed using Student *t* test. Significance was assumed when *p* < 0.05. \**p* < 0.05, \*\**p* < 0.01, \*\*\**p* < 0.001, \*\*\*\**p* < 0.0001.

We further measured changes in myelin content of CA3 Shaffer collateral axons, the axons projecting from CA3 pyramidal cells to CA1. Using G-ratio analysis, an inverse indicator of myelin content of an axon fiber (Egawa et al., 2018)(Fig. 6F), we found a significant increase in the G-ratio of AD-*SynRFP* mice compared to WT-*SynRFP* at both 9 and 11 m (Fig. 6G), while AD-*SynCav1* mice exhibited a significant decrease in G-ratio versus AD-*SynRFP*. Further analysis revealed that the G-ratio change was due to altered myelin sheath thickness (Fig. 6H) rather than axon lumen diameter (Fig. 6I).

### *SynCav1* preserves ultrastructural indicators of synaptic plasticity in AD mice

Loss of synapses in AD patients is closely linked to cognitive deficits and dementia(Honig et al., 2018; Hyman et al., 2012; Scheff, Price, Schmitt, DeKosky, & Mufson, 2007; Selkoe, 1991). We next assessed synaptic ultrastructure of CA1 distal apical dendrites in the *stratum radiatum* using EM. Hippocampal CA1 distal apical dendrites (Fig. 7A) exhibited decreased total type I excitatory asymmetric synapses (Fig. 7B) and total number of PSVs/bouton (Fig. 7C) in AD-*SynRFP* mice versus WT-*SynRFP*, findings that consistent with Scheff et al. which demonstrated decreased synapses in the *stratum radiatum* within the CA1 subfield in AD patients(Scheff et al., 2007). When compared to AD-*SynRFP*, AD-*SynCav1* mice exhibited preserved total synapses and PSVs/bouton (Fig. 7B, C). In addition, AD-*SynCav1* exhibited more total PSVs/bouton versus WT-*SynRFP* at 9 m. As shown in Fig. 7D, we also observed altered dendritic spine morphology (i.e., more-stubby and less mushroom-like spines) in AD-*SynRFP* mice compare to WT-*SynRFP*, a finding similar to that observed in human AD biopsies and AD transgenic models(Androuin et al., 2018). In contrast, while AD-*SynCav1* mice showed preserved neck diameter) (Fig. 7E) and increased dendritic spine length (Fig. 7F) compared to AD-*SynRFP* mice.

**Fig. 7.**
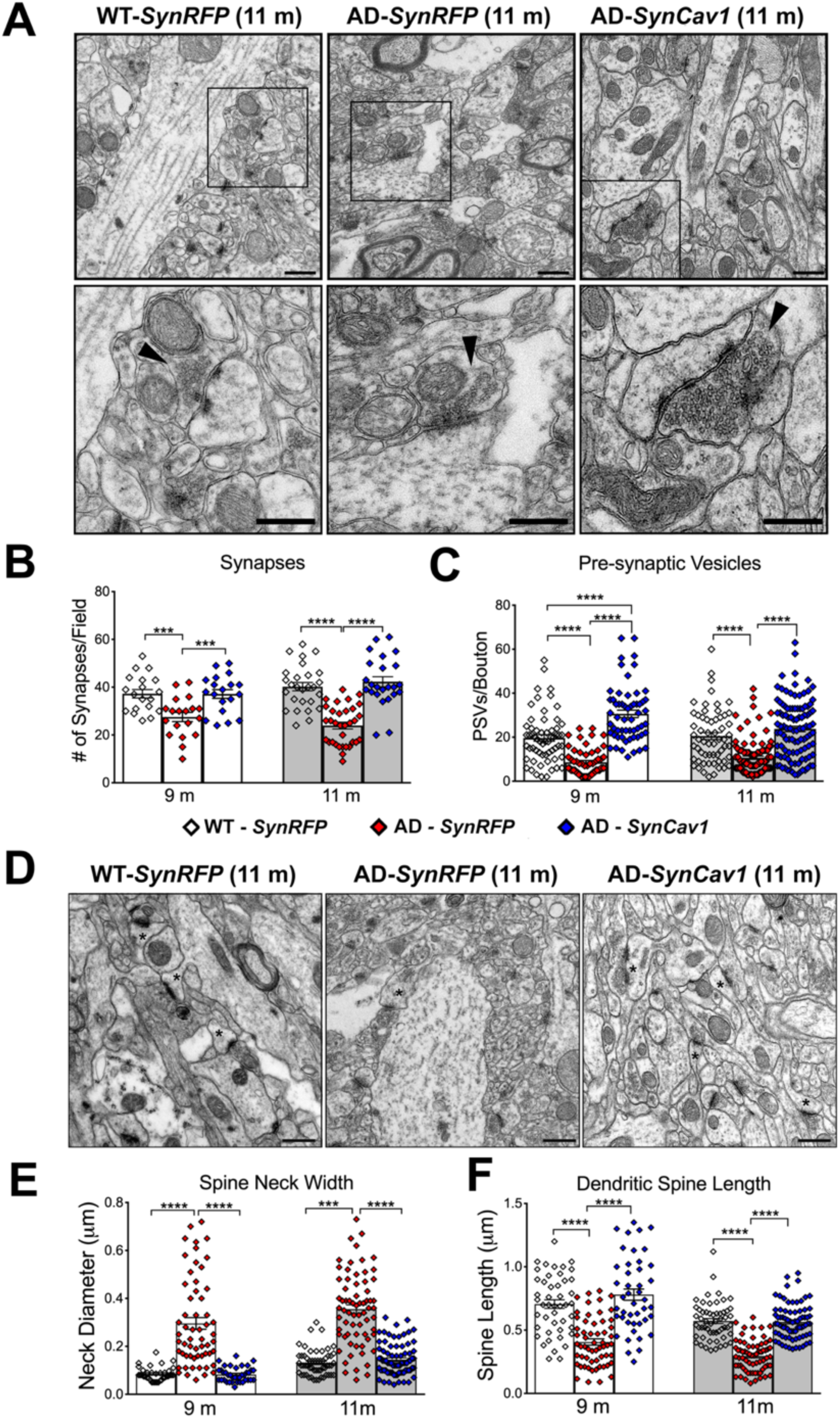
*SynCav1* gene delivery preserves synaptic ultrastructure and dendritic spine morphology in the hippocampus of 9 m and 11 m AD mice. EM images (**A**) and quantitation of total type I asymmetric synapses (**B**) and presynaptic vesicles (PSVs)/axonal bouton (**C**) in hippocampal CA1 distal apical dendrites in the *stratum radiatum.* AD-*SynRFP* mice demonstrated a significant reduction in total type I excitatory asymmetric synapses and total number of pre-synaptic vesicles (PSVs)/bouton. In contrast, AD-*SynCav1* mice showed a significant increase in both total synapses and total number of PSVs/bouton when compared to AD-*SynRFP* and WT-*SynRFP*. **Top micrographs** represent 4800x Magnification, scale bar = 1 um; **bottom micrographs** represent 11000x Magnification, scale bar = 500 nm. Black arrow heads denote asymmetric type I synapses. EM images (**D**) and quantitation of dendritic spine neck width (**E**), length (**F**) showed that AD-*SynRFP* mice exhibited altered dendritic spine morphology (i.e., more-stubby and less mushroom-like spines) with significantly increased neck diameter and reduced spine length compare to WT-*SynRFP*. AD-*SynCav1* mice showed preserved neck diameter and increased spine length when compared to AD-*SynRFP*. **Same trend was observed** at 11month age. Micrographs represents 11000x Magnification, scale bar = 500 nm. Data are presented as mean ± SEM, *n* = 3-4 animals per group consisting of 10 electron micrographs per animal. Data were analyzed using Student *t* test. Significance was assumed when *p* < 0.05. \**p* < 0.05, \*\**p* < 0.01, \*\*\**p* < 0.001, \*\*\*\**p* < 0.0001.

### *SynCav1* does not mitigate amyloid plaque deposition or gliosis in 11 m AD mice

To assess whether *SynCav1* had an effect on amyloid plaque deposition and astrogliosis in PSAPP mice, we performed IF on brain tissue at 11 m using antibodies to 6E10. IF showed similar levels of hippocampal 6E10-positive amyloid plaques deposition in AD-*SynRFP* and AD-*SynCav1* mice (Fig. 8A). Co-localization between Cav-1 and 6E10-positive plaques (Fig. 8C) was detected in 11 AD brain tissue. To assess whether *SynCav1* had an effect on astrogliosis in PSAPP mice, we performed IF on brain tissue at 11 m using antibodies to GFAP. Both AD-*SynRFP* and AD-*SynCav1* brains also exhibited elevated GFAP expression (Fig. 8D,E), indicating that *SynCav1* did not mitigate amyloid plaque deposition or astrogliosis in 11 m old PSAPP mice.

**Fig. 8.**
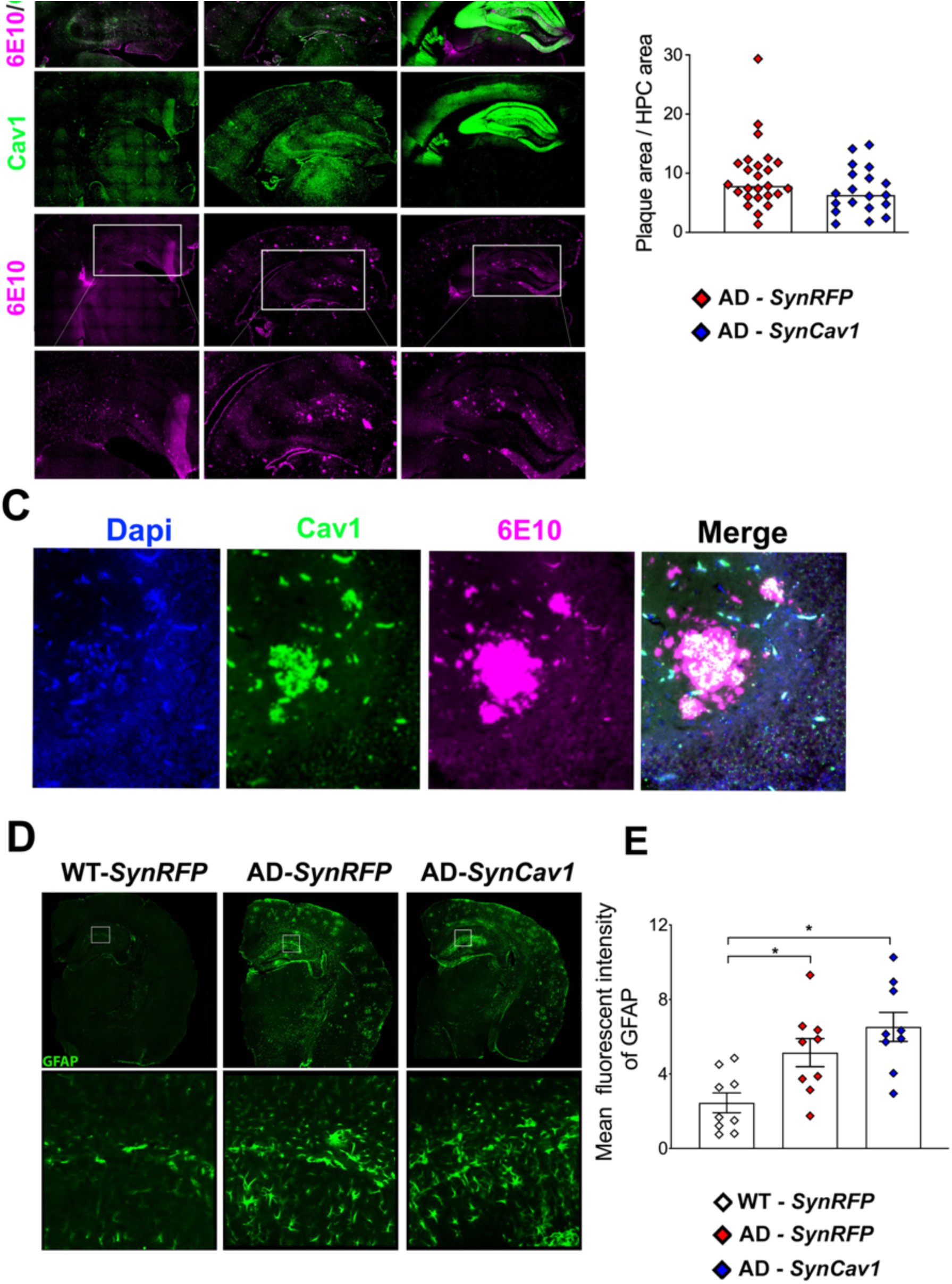
*SynCav1* did not mitigate hippocampal amyloid plaque deposits nor astrogliosis in 11 m AD mice. Cav1/6E10 dual staining of 11month AD mouse hippocampus (**A**) and 6E10-positive amyloid deposits quantitation (**B**) in WT-*SynRFP*, AD-*SynRFP*, and AD-*SynCav1* hippocampus at 11 m. (**C**) Cav1/6E10 dual staining of 11month AD mouse hippocampus sub-region showed a colocalization of Cav1 with amyloid plaque, cell nuclei were counterstained with DAPI (blue). GFAP (astrocytes) (**D**) with quantitation (**E**) in WT-*SynRFP*, AD-*SynRFP*, and AD-*SynCav1* hippocampus at 11 m. Data were analyzed using Student *t* test, *n* = 3-4 animals per group. Significance was assumed when \**p* < 0.05.

## Discussion

The present study is the first to demonstrate that a one-time *SynCav1* gene delivery preserves hippocampal structure and hippocampal-dependent cognitive function in an AD mouse model (*PSAPP*). Furthermore, *SynCav1* preserved MLR-localization of protein markers of neuronal and synaptic plasticity, all of which occurred independent of removing neurotoxic Ab plaques or reducing astrogliosis. This study builds upon previous work demonstrating the neuroprotective effects of *SynCav1* gene therapy in the setting of aging, traumatic brain injury, and ALS(Egawa et al., 2017; Mandyam et al., 2017; Sawada et al., 2019), and now further extends the therapeutic potential of *SynCav1* in another neurodegenerative disease indication, i.e., the *PSAPP* transgenic model of AD.

Individuals with severe AD exhibit decreased neurotrophic signaling in the cortex and hippocampus(Ferrer et al., 1999), suggesting that delivery of neurotrophins(Tuszynski et al., 2005) or increasing neurotrophin receptor (NTR) signaling could potentially reverse cognitive deficits in AD(Beeri & Sonnen, 2016; Castello et al., 2014). However, much evidence shows that functional neurotrophin signaling is dependent upon intact MLRs, Cav-1 and NTR localization to MLRs(Head et al., 2011; Mandyam et al., 2017). Pre-clinical models demonstrated that toxic Ab or abnormal tau exposure decreased Cav-1 protein expression in neurons (Wu et al., 2017; Yuan et al., 2017), while interventions that restored or augmented Cav-1 mitigated abnormal tau accumulation, Ab production, and reversed neurotoxicity(Kapoor et al., 2010; Wu et al., 2017; Yuan et al., 2017; Y. L. Zhao et al., 2016), suggesting that Cav-1 restoration may preserve MLR-associated neuroprotective synaptic signaling complexes, thus affording neuronal resilience against toxic Ab and abnormal tau. The present study extends the knowledge that augmenting Cav-1 specifically in neurons increases full-length TrkB (fl-TrkB) expression in MLRs which can undergo autophosphorylation upon agonism to activate intracellular signaling pathways necessary for neurite outgrowth, dendritic spine morphology, and cognitive performance(Fenner, 2012; Psotta et al., 2015). Considering that we did not observe significant difference in fl-TrkB expression in whole tissue hippocampal homogenates between WT the AD mice, the decreased fl-TrkB in MLRs in AD mice suggests disrupted subcellular localization of fl-Trkb attributed to either disrupted MLRs or decreased Cav-1 expression within MLRs. This failure in fl-TrkB localization from non-MLR subcellular regions to buoyant MLRs may explain the lack of efficacy of exogenous neurotrophin delivery in several failed clinical trials. The current study suggests that preservation of MLR-associated fl-TrkB expression in AD-*SynCav1* mice may potentially increase the efficacy of endogenous (BDNF) or exogenous agonists (e.g., dihydroxyflavone(Castello et al., 2014; Chen et al., 2018; English, Liu, Nicolini, Mulligan, & Ye, 2013)) that specifically target TrkB activation.

AD brains exhibit anatomical and ultrastructural changes (i.e., reduced synapses) in hippocampal and cortical brain regions(Liu, Erikson, & Brun, 1996). Although deposition of amyloid plaques is the traditional hallmark of AD, decreased synapses, “mushroom” spines (i.e., synaptopathy) and neuronal atrophy more closely correlate with cognitive impairment(Androuin et al., 2018; Price et al., 2001; Terry et al., 1991). We previously demonstrated that *SynCav1* enhanced hippocampal neuroplasticity and improved cognitive function in adult and aged WT mice(Egawa et al., 2018; Mandyam et al., 2017). The present study further extends the neuroprotective capacity of *SynCav1* gene therapy in a neurodegenerative AD mouse model, independent of attenuating toxic amyloid deposition or astrogliosis. Furthermore, *SynCav1* preserved synaptic ultrastructure, dendritic spine morphology, and axonal fiber myelin content in these AD mice. Noticeable, although AD-*SynRFP* mice exhibit less synapses at 9 m, increased PSD length and area were observed (data not shown), a subtle compensatory ultrastructural change consistent with Scheff et al. and others(DeKosky & Scheff, 1990; Scheff, Sparks, & Price, 1996), which may explain for the normal contextual memory observed at 9 m in AD mice.

MLRs are specialized plasmalemmal microdomains that compartmentalize and modulate synaptic signaling and adaptor proteins essential for synaptogenesis, synaptic plasticity and cognitive function(Egawa, Pearn, Lemkuil, Patel, & Head, 2016; Egawa et al., 2018; Mandyam et al., 2017). Proteomics on MLR fractions revealed a novel association between hippocampal MLRs and the AMPAR adaptor and regulatory protein Shisa9 (Karataeva et al., 2014; Khodosevich et al., 2014; Kunde, Rademacher, Zieger, & Shoichet, 2017). Shisa9 is known to regulate synaptic plasticity and memory formation in the hippocampus(Karataeva et al., 2014; Khodosevich et al., 2014). Proteomics revealed that MLR-associated Shisa9 was significantly decreased in AD mice, yet preserved with *SynCav1*, suggesting that Shisa9 may contribute to the neuroprotective phenotype in *SynCav1*-treated AD mice. Immunoprecipitation of MLR fractions using Cav-1 antibodies confirmed the colocalization of Cav-1 with Shisa9 in hippocampal tissue (Supplementary Fig.4). The preservation of MLR-associated Shisa9 in the AD-*SynCav1* hippocampus, may in part explain for the neuroanatomical preservation of mossy fiber projections from the dentate gyrus to CA3 hippocampal subfield in AD-*SynCav1* mice. We are currently researching whether *SynCav1*’s neuroprotective effect in the present study is dependent upon Shisa9/AMPAR signaling in MLRs. Additional, proteomic analyses revealed reduced expression of several neurodegenerative-related pathways in MLRs from AD-*SynCav1* mice, suggesting the importance of plasma membrane microdomains in modulating protein complexes involved in neuronal function and resilience against neurodegenerative conditions.

There are conflicting results regarding Cav-1 expression levels in the AD brain; while the present study and others have shown decreased Cav-1 protein and mRNA in the AD and neurodegenerative brain(Mufson et al., 2018), Gaudreault et al. detected increased Cav-1 protein and mRNA in the hippocampus and frontal cortex of AD brains(Gaudreault, Dea, & Poirier, 2004). It is well known that both astrocytes and microglia express Cav proteins(Ikezu et al., 1998; Niesman et al., 2013; Takeuchi, Matsuda, Tooyama, & Yasuhara, 2013). Thus, the elevated Cav-1 observed by Gaudreault et al may due to prominent astrogliosis and microgliosis in AD brains(Glass, Saijo, Winner, Marchetto, & Gage, 2010). Activated microglia play a central role in clearing debris such as amyloid plaques(R. Zhao, Hu, Tsai, Li, & Gan, 2017). The present study showed strong co-localization between Cav-1 and 6E10-positive plaques at 11 m (Fig. 8C), which may explain the abnormal increase of Cav-1 in 11month AD mice (data not shown here). More study is needed to understand the source of this increased Cav-1. In a recent study using postmortem human neuropathological brain samples, Mufson et al. revealed decreased Cav-1 mRNA level specifically in degenerating neurons while non-degenerating neurons still had normal Cav-1 level. The present study detected decreased hippocampal Cav-1 expression in AD mice as earlier as 6 months with an associated fear learning deficit (data not shown). Considering that 9 month AD mice exhibited decreased hippocampal Cav-1 expression along with learning and memory deficit, it is plausible that loss of Cav-1 expression in the hippocampus may contribute to neurodegeneration(Head et al., 2010; Mufson et al., 2018; Niesman et al., 2014). While the current study provides proof-of-concept for the neuroprotective and therapeutic potential of *SynCav-1* gene therapy in AD, we are also currently testing whether *SynCav1* delivery to PSAPP mice at 6 months of age can reverse learning and memory deficits.

In summary, the present study demonstrates that *SynCav1* gene delivery delays neurodegeneration and cognitive deficits, preserves hippocampal arborization, synaptic integrity, and axonal ultrastructure in AD mice. Furthermore, *SynCav1* preserves MLR-localization of the synaptic components essential for neuronal and synaptic plasticity independent of attenuating toxic Ab plaque accumulation or astrogliosis. Therefore, *SynCav1* may serve as a novel gene therapy to preserve or delay neurodegenerative conditions in AD and other forms of CNS disease of unknown etiology.

## Methods and Materials

### Animals

AD-transgenic (Tg) (*APPSwePS1d9* a.k.a. *PSAPP*, Supplementary Fig. 1) and C57BL/6 mice were purchased from Jackson Laboratory (Bar Harbor, ME, USA) and bred in-house. Wild type (WT) littermates were used as control. All animal protocols were approved by the Veterans Administration San Diego Healthcare System Institutional Animal Care and Use Committee (#14-044). Mice were reared (3-5/cage) with free access to food and water. AAV9-*SynRFP* was used as the control vector, and neuronal targeted overexpression of Cav-1 was achieved by AAV9-*SynCav1* viral vector (Fig. 1A). At 3 months (m) of age, WT and AD mice were allocated to 3 groups randomly: WT-*SynRFP*, AD-*SynRFP* and AD-*SynCav1* to receive hippocampal stereotactic injections. Brain tissue was processed for biochemistry, histopathology, immunofluorescence and electron microscopy after behavior tests (Fig.1C).

### Stereotactic Injection

Three-month-old mice were mounted onto a stereotaxic frame under anesthesia (2% isoflurane). Bilateral burr holes were made by 22-gauge needle. Hippocampal injections using 33-gauge, 10-µL Gas Tight syringe (Hamilton, Reno, Nevada) were controlled by injectomate (Neurostar, Berlin, Germany). 1.5 µL of adeno-associated virus serotype 9 (AAV9) (viral titer: 10^9^ genome copies (g.c.)/µL) containing synapsin-red fluorescent protein or synapsin-caveolin-1 (*SynCav1*) was injected bilaterally over 180 seconds at three locations (1st site: AP: 1.82 mm, Lat: 1.15 mm, DV: 1.7 mm: 2nd site: AP: 2.30 mm, Lat: 2.25 mm, DV: 1.75 mm; 3rd site: AP: 2.80, Lat: 2.5 mm, DV: 2.00 mm) with 1 minute indwelling time as previously described(Mandyam et al., 2017). Fig. 1B and Supplemental Video. 2 display broad brain tissue AAV infectivity and transgene expression (whole hippocampus rostral to caudal, dorsal to ventral and portions of the somatosensory and parietal cortex) using *SynRFP.*

### Open Field and Fear Conditioning Behavior

Open field and fear conditioning were performed as previously described(Mandyam et al., 2017). Locomotion was recorded for 10 min and analyzed by a computerized video tracking system (Noldus XT 7.1, Leesburg, VA). Recorded parameters included distance moved (cm), velocity (cm/second) and time spent in the center of the arena (seconds). For fear conditioning, presentation of unconditioned stimuli (US) (foot-shock) and conditioned stimuli (CS) (auditory tone) were controlled using Med Associates Inc. (St. Albans, Vermont), and movement was monitored by video. Freezing protocol (Fig. 1D) was determined using Video Freeze (Med Associates Inc.; San Diego Instruments, San Diego, California)(Mandyam et al., 2017; Pearn et al., 2018).

### Biochemical Characterization of Membrane/Lipid Rafts (MLRs)

Hippocampi were homogenized at 4°C in 500 mM sodium carbonate (pH 11.0) (containing protease and phosphatase inhibitors) and then sonicated 3x for 15 s. Samples (0.5 mg/ml) were subjected to sucrose density gradient fractionation as previously described(Mandyam et al., 2017). Homogenates and fractions were immunoblotted using primary antibodies Cav-1 (Cell Signaling #3238; 1:1000), TrkB (BD Biosciences 610102; 1:1000), synaptobrevin (BioLegend #837001, 1:1000) and GAPDH (Cell Signaling #2118s; 1:1000) overnight at 4°C followed by incubation with IR-dye labeled secondary antibody for 1 hour and measured with LiCor Odyssey followed by densitometric analysis.

### Proteomics

MLRs protein samples were diluted in TNE (50 mM Tris pH 8.0, 100 mM NaCl, 1 mM EDTA) buffer. RapiGest SF reagent (Waters Corp.) was added to the mix (0.1% final concentration) and samples boiled for 5 minutes. 1 mM TCEP (Tris (2-carboxyethyl) phosphine) was added to the samples and incubated at 37°C for 30 minutes. Samples were carboxymethylated with 0.5 mg/ml of iodoacetamide for 30 minutes at 37°C followed by neutralization with 2 mM TCEP. Samples were digested with trypsin (trypsin: protein ratio - 1:50) overnight at 37°C. RapiGest was degraded and removed by treating the samples with 250 mM HCl at 37°C for 1 hour followed by centrifugation at 14k r.p.m. for 30 minutes at 4°C. The soluble fraction was then added to a new tube and the peptides were extracted and desalted using C18 desalting columns (Thermo Scientific, PI-87782). Peptides were quantified using BCA assay and a total of 1 ug of peptides were injected for LC-MS analysis.

### LC-MS-MS

Trypsin-digested peptides were analyzed by ultrahigh pressure liquid chromatography (UPLC) coupled with tandem mass spectroscopy (LC-MS/MS) using nano-spray ionization using an Orbitrap fusion Lumos hybrid mass spectrometer (Thermo) interfaced with nano-scale reversed-phase UPLC (Thermo Dionex UltiMate™ 3000 RSLC nano System) using a 25 cm, 75-micron ID glass capillary packed with 1.7-µm C18 (130) BEHTM beads (Waters corporation). Protein identification and label free quantification was carried out using Peaks Studio 8.5 (Bioinformatics solutions Inc.). Proteins were considered identified if false discovery rate (FDR) was less than 1% of peptide-spectrum matches to *m. musculus* database (Uniprot/SwissProt) compared to decoy-fusion library.

### Bioinformatics

Quantifiable protein levels were expressed as peak area and compared as relative intensity for WT-*SynRFP* versus AD-*SynRFP* or AD-*SynRFP* versus AD-*SynCav1* with significance set at *p* < 0.05. Mean significantly different expression levels were displayed as heatmaps for up- or down-regulation using Euclidean clustering with complete linkage. Significantly up or down-regulated quantifiable protein groups were subjected to ontology (GO) using STRING-db (www.string-db.org). The GO terms with lowest FDR (all less than 1%) were displayed. Significantly regulated GO terms are displayed as log_2_ fold change of genes.

### Golgi-Cox Staining

Brains were submerged in Golgi-Cox solution A + B (FD Neurotechnologies Inc., Ellicot City, Maryland) for 8 days followed by solution C for 4 days at room temperature. 80-µm-thick coronal cryo-sections were prepared for staining with solution D + E and dehydrated according to the manufacturer’s instructions. Neurolucida (Micro-BrightField, Willston, Vermont) and NeuroExplorer were used to trace and analyze dendritic branching (15 neurons from 3 mice/group)(Mandyam et al., 2017).

### Immunofluorescence Microscopy

Floating sections were blocked with 10% goat serum and incubated with chicken anti-MAP-2 (1:250, Abcam, USA) and rabbit anti-Cav-1 (1:1000, Cell Signaling, USA) at 4°C overnight. Slices were then incubated with species-specific fluorescence secondary antibody in the dark for 1 hour and preserved with anti-fade Dapi-mounting medium. For amyloid-β plaques staining, sections were incubated in 98% formic acid for 5 min to expose the epitope prior to the blocking process, and purified Alex fluor 488 anti-β-Amyloid 1-16 antibody (1:200, BioLegend) was used to visualize amyloid plaques.

### Electron Microscopy

Electron microscopy was performed as previously described(Egawa et al., 2018). Excitatory synapses were identified by the presence of a prominent, asymmetric postsynaptic density (PSD), and pre-synaptic vesicles (PSVs) were normalized to the total boutons per field. Morphological dendritic spines were measured as previously described(Androuin et al., 2018). G-ratio was defined as the diameter of the axon lumen divided by the diameter of the fiber (axon lumen plus myelin)(Egawa et al., 2018).

### Statistical Analyses

Data were checked for normal distribution, all behavior data were analyzed by one-way analysis of variance (ANOVA) or 2-way ANOVA followed by Fisher’s LSD or Tukey’s multiple comparisons tests and all biochemical data were analyzed by Student *t* tests as appropriate using GraphPad Prism 6 (La Jolla, California). Data were presented as mean ± SEM and significance assumed when \**p* < 0.05. Experimental groups were blinded to the observer and code was broken for analysis.

## Acknowledgements and Disclosures

This manuscript is dedicated to the memory of Patrick J. Head and Marilyn G. Farquhar. Thanks to Alice Zemljic-Harpf (Anesthesiology) and Ying Jones (UCSD EM core) for preparation of the brain samples and training on the EM and Dr. Marilyn Farquhar for consultation. Electron micrographs were taken in the Cellular and Molecular Medicine EM core facility supported in part by National Institutes of Health Award number S10OD023527. Thanks to Robert Rissman, Director of UCSD Alzheimer’s Disease Cooperative Study (ADCS) Biomarker Core and AD Research Center (ADRC) Neuropathology Core, for his technical assistant with amyloid plaque immunofluorescence using 6E10 antibody. Thanks to Chitra Mandyam for consultation on behavior. Work in the authors’ laboratories is supported by Veterans Affairs Merit Award from the Department of Veterans Affairs BX003671 (B. P. Head); National Institutes of Health, Bethesda, MD, U.S.A., NS073653 (B. P. Head), GM085179 (P. M. Patel), R56AG057469 (V. Hook), and R01NS109075 (V. Hook). J.Q.T. was supported by the Penn AD Core Center grant (AG-10124-28). S. Wang, J.S. Leem, S. Podvin, V. Hook, N. Kleschevnikov, P. Savchenko, M. Dhanani, K.Y.Q. Zhou, I.C. Kelly, T. Zhang, A. Miyanohara, A. Kleschevnikov, S.L. Wagner, J.Q. Trojanowski, D. M. Roth, and P. M. Patel having no financial disclosures. H.H. Patel and B.P. Head are scientific founders of CavoGene LifeSciences LLC.

## Author Contributions

S.W. performed experiments, data analyses, wrote and edited the manuscript; J.S.L. performed genotyping, colony maintenance and biochemical experiments; S.P. and V.H. assisted with proteomic analyses and bioinformatics; N.K. performed behavioral studies; P.S. performed biochemical experiments; M.D. performed biochemical experiments; K.Y.Q.Z. assisted with vector gene delivery; I.C.K. assisted with Golgi-Cox analyses; T.Z. performed the light sheet microscopy; A.M. generated the AAV vector constructs; A.K. assisted with experimental design; S.L.W. assisted with experimental design and editing; J.Q.T. assisted with microscopic experimental design and editing; D.M.R., H.H.P., and P.M.P. assisted with editing; B.P.H. supervised overall design and concept, performed data analyses, wrote and edited the manuscript.

## Supplementary Materials

### Materials and Methods

#### Genotyping

*APPSwePS1d9* mice were confirmed by genomic DNA extraction and PCR using the Qiagen DNeasy Blood and Tissue Kit (69504; Qiagen, Valencia, CA, USA). PCR was performed for *APP* genes by using the following protocol: denaturation at 94°C for 2 min, followed by 10 cycles at 94°C for 20 s, 65°C for 15 s (−0.5 per cycle decrease), and 68°C for 10 s, then followed by 28 cycles at 94°C for 15 s, 60°C for 15 s, 72°C for 10 s, then 72°C for 2 min and hold at 10°C. All primers were purchased from Integrated DNA Technologies (Coralville, IA, USA). Primer sequences to detected transgenic positive (400 bp and 750 bp): Prp-Sense-J (mouse prion protein variant 1-2 mRNA) 5′-GGG ACT ATG TGG ACT GAT GTC GG -3′, Prp-Antisense J (mouse prion protein variant 1-2 mRNA), 5′-CCA AGC CTA GAC CAC GAG AAT GC -3′, and S36 mouse amyloid precursor protein (APP) mRNA), 5’-CCG AGA TCT CTG AAG TGA AGA TGG ATG -3’. Transgenic positives were identified by two bands at 400 and 750 bp, while transgene negative presented 750 bp only (Figure S1). PCR products were separated on a 1% agarose gel (35 min at 135 volts).

**Supplementary Fig. 1.**
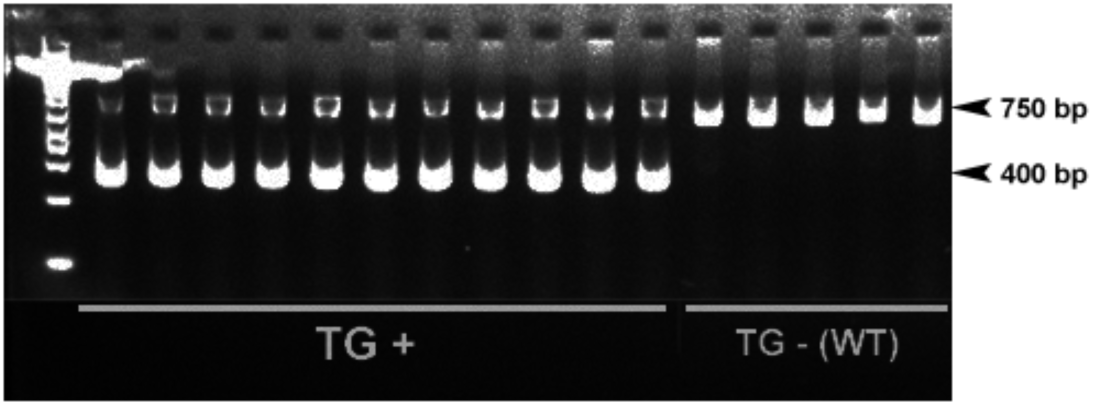
PCR and gel electrophoresis confirm positive double transgenic mice expressing a chimeric mouse/human amyloid precursor protein (Mo/HuAPP695swe) and a mutant human presenilin 1 (PS1-dE9).

**Supplementary Video. 2** Light Sheet Microscopy AAV9-*SynRFP* injected mouse hippocampus (1-week post-injection), optically cleared in X-CLARITY hydrogel solution (Cat #13103). Data processing and 3D rendering was done using Arivis Vision4D™. Scale bar = 100 um

**Supplementary Fig. 3.**
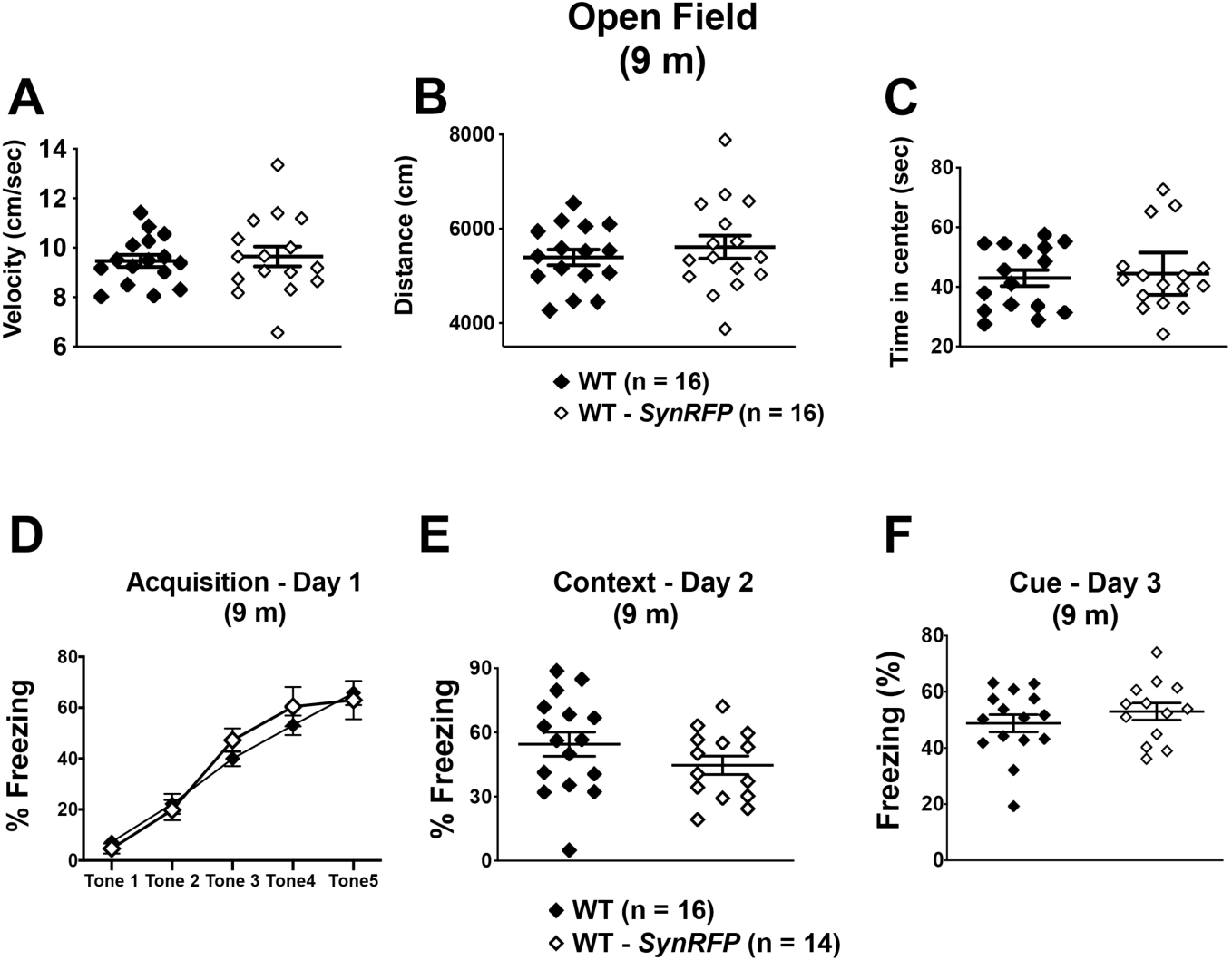
Open field and fear conditioning test comparing naïve WT with WT-*SynRFP* at 9 m. (**A-C**) Open field (velocity, distance moved, time in center) and (**D-F**) fear conditioning for naïve WT (*n* = 16) versus WT-*SynRFP* (*n* =14-16) at 9 m. For open field data (mean ± SEM) and fear conditioning data (percent (%) freezing mean ± SEM) were analyzed using Student *t* test (Day 2 and 3). Significance was assumed when *p* < 0.05.

**Supplementary Fig. 4.**
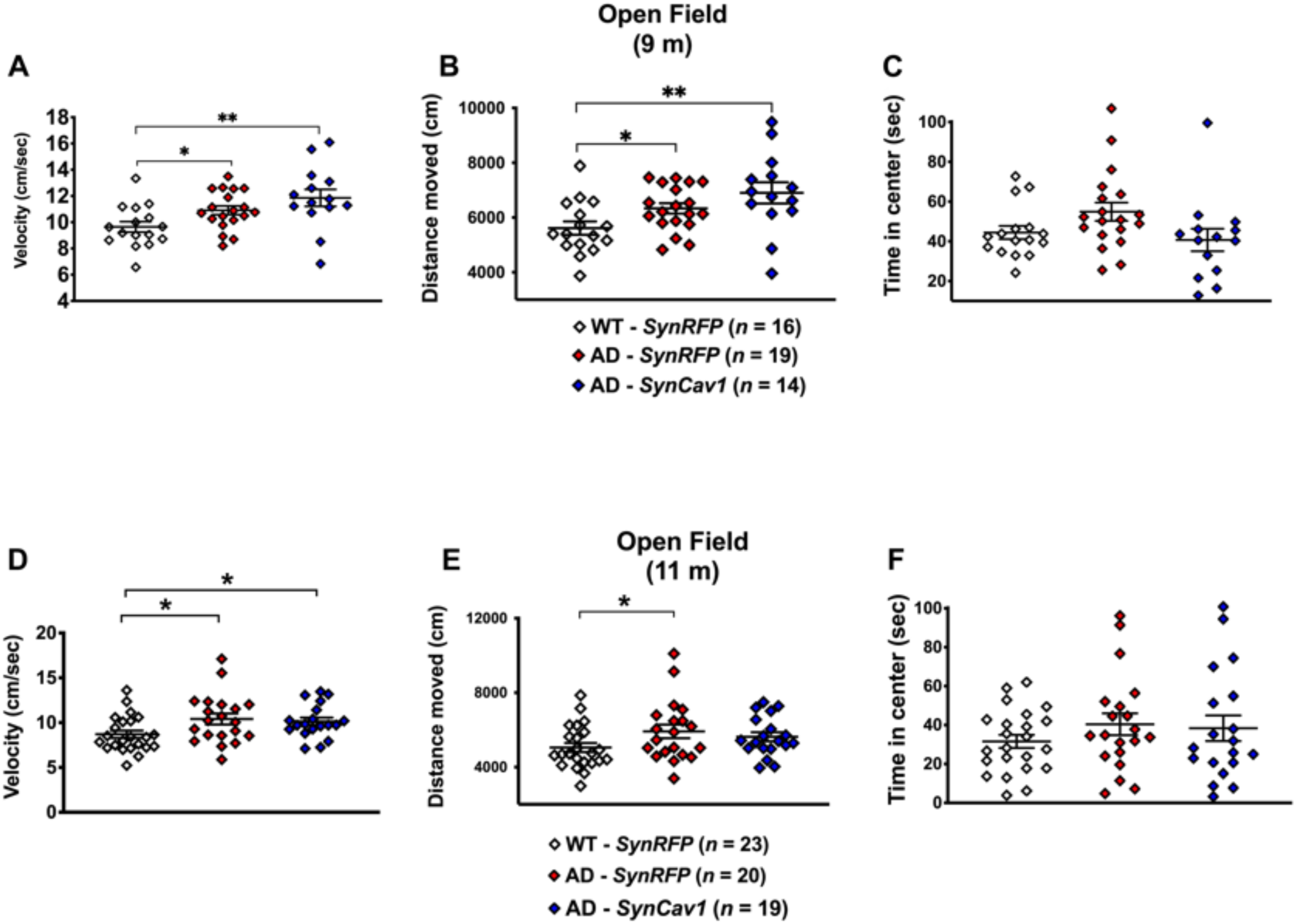
Open field performance of 9 and 11 m *APPSwePS1d9* mice. (A, **D**) Mean velocity, (**B, E**) distance traveled, and (**C, F**) time spent in the center of 9 and 11-month old WT-*SynRFP* (*n* = 16 at 9 m; *n* = 23 at 11 m), AD-*SynRFP* (*n* = 19 at 9 m; *n* = 20 at 11 m), and AD-*SynCav1* (*n* = 14 at 9 m; *n* = 19 at 11 m) mice respectively. For open field data (mean ± SEM) was analyzed using Student *t* test. Data are presented as mean ± SEM. Significance was assumed when *p* < 0.05. \**p* < 0.05, \*\**p* < 0.01.

**Supplementary Fig. 5.**
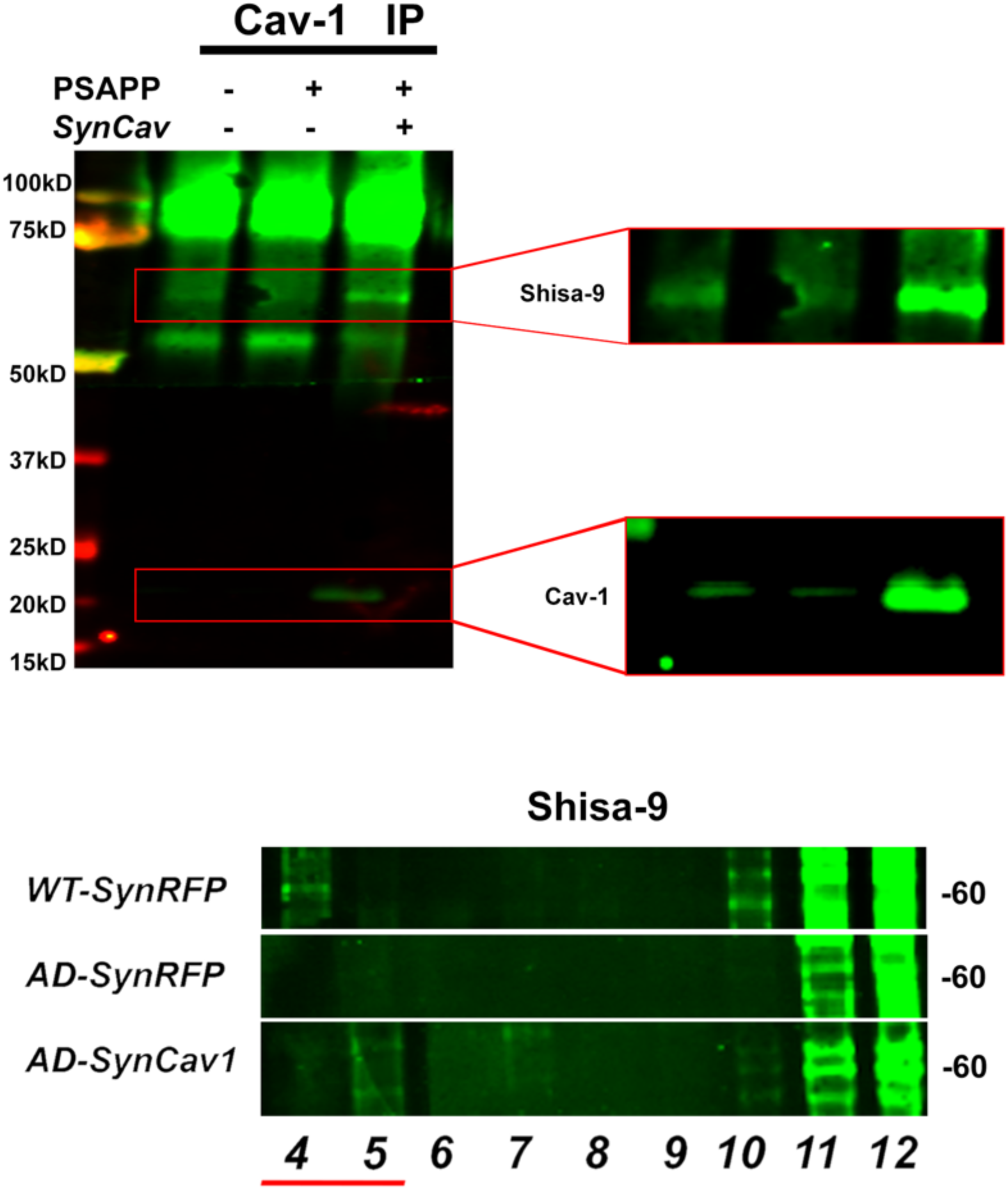
Cav-1 IPs of MLR fractions confirm the interaction of Shisa9 and Cav1 in mice hippocampi **(A)**. Cav-1 IP of MLR fractions revealed decreased Shisa9 expression in *AD-SynRFP* and the highest Shisa9 expression in AD-*SynCav1*. (B) Representative blot of Shisa9 in hippocampal MLR fractions in WT-SynRFP, AD-SynRFP, and AD-SynCav1 mice at 9 m.

